# Development and Application of Cas13a-based Diagnostic Assay for *Neisseria Gonorrhoeae* Detection and Identification of Azithromycin Resistance

**DOI:** 10.1101/2021.05.20.445076

**Authors:** Hao Luo, Wentao Chen, Zhida Mai, Xiaomian Lin, Jianjiang Yang, Lihong Zeng, Yuying Pan, Qinghui Xie, Qingqing Xu, Xiaoxiao Li, Yiwen Liao, Zhanqin Feng, Jiangli Ou, Xiaolin Qin, Heping Zheng

## Abstract

Gonorrhea caused by *Neisseria gonorrhoeae* has spread world-wide. Antimicrobial-resistant strains have emerged to an alarming level to most antibiotics including to the ceftriaxone-azithromycin combination, currently recommended as first-line dual therapy. Rapid testing for antimicrobial resistance will contribute to clinical decision-making for rational drug use and will slow this trend. Herein, we developed a Cas13a-based assay for *N. gonorrhoeae* detection (porA target) and azithromycin resistance identification (A2059G and C2611T point mutations). We evaluated the sensitivity and specificity of this method, and 10 copies per reaction can be achieved in porA detection and C2611T identification, with no cross-reactions. Comparison of the Cas13a-based assay (porA target) with Roche Cobas 4800 assay (n=23 urine samples) revealed 100% concordance. Isolated *N. gonorrhoeae* strains were used to validate the identification of A2059G and C2611T resistance mutations. All tested strains (8 A2059G strains, 8 C2611T strains, and 8 wild-type strains) were successfully distinguished by our assay and verified by testing MIC for azithromycin and sequencing the 23S rRNA gene. We adopted lateral flow for the SHERLOCK assay readout, which showed a visible difference between test group and NC group results. To further evaluate the capability of our assay, we tested 27 urethral swabs from patients with urethritis for *N. gonorrhoeae* detection and azithromycin-resistance identification. Of these, 62.96% (17/27) strains were detected with no mutant strains and confirmed by sequencing. In conclusion, the novel Cas13a-based assay for rapid and accurate *N. gonorrhoeae* detection combined with azithromycin drug resistance testing is a promising assay for application in clinical practice.

## Introduction

Gonorrhea is a common bacterial sexually transmitted infection (STI) in the world, caused by *Neisseria gonorrhoeae* [1]. As estimated by the World Health Organization (WHO), there were 78 million global cases in 2012 and 86.9 million cases in 2016 worldwide [2, 3]. The prevalence of *N. gonorrhoeae* has increased rapidly and remains a public health concern. In the absence of an effective vaccine, antibiotic treatment is critical to cure and slow the spread of *N. gonorrhoeae* infections [4, 5]. However, due to the use and abuse of antibiotics, antimicrobial resistance (AMR) of *N. gonorrhoeae* has emerged to all first-line therapeutic drugs used to date [4–7]. In particular, AMR to azithromycin and ceftriaxone currently used as first-line dual therapy has been reported as a cause of treatment failure in both the United Kingdom and Australia [8, 9], and the resistance has shown a gradual increasing trend according to the gonococcal surveillance program data from Europe and the United States [10, 11]. There is, therefore, a need for clinicians to rapidly acquire resistance data for antibiotics, which could help manage rational drug use and further slow the development of drug resistance.

Traditional antimicrobial resistance detection methods are mainly culture-based. The quantitative agar dilution method which can determine the minimum inhibitory concentration (MIC) of antimicrobials is recognized as the ‘gold standard’ method, but complicated protocol steps and long turnaround times hinder its development to satisfy clinical requirements [1, 6]. To achieve this goal, non-culture-based nucleic acid amplification tests (NAATs) have been introduced and are being developed. Sequencing technology has been widely used to identify plasmid-mediated or chromosomally-mediated drug resistance to discover antimicrobial resistance towards penicillin, ciprofloxacin, tetracycline, azithromycin, extended-spectrum cephalosporin, and multidrug resistance [6, 12]. With the discovery of a strong correlation between single nucleotide polymorphisms (SNP) and drug resistance in *N. gonorrhoeae*, more convenient assays have been established [13–17]. These methods generally use PCR to amplify target genes, and combine with specific probes, high resolution melting (HRM) analysis, or mass spectrometry to differentiate specific point mutations. The protocol is time-saving, however, large precision instruments are necessary to ensure accurate temperature control and results also require skilled evaluation.

Cas13a was first described by Zhang et al. and exploits endonuclease activity of target RNA and collateral cleavage activity of the target sequence [18]. Based on this principle, the SHERLOCK (<u>s</u>pecific <u>h</u>igh-sensitivity <u>e</u>nzymatic <u>r</u>eporter un<u>lock</u>ing) was developed, which combines recombinase polymerase amplification (RPA) and Cas13a in an isothermal system with single molecule sensitivity, high specificity, single-base resolution, and convenient acquisition of results [19]. Given the advantages of this technology, SHERLOCK is becoming a potential tool for rapid diagnostic testing of emerging infectious diseases, and has been developed for the diagnosis of *Plasmodium*, SARS-CoV-2, Ebola virus, and Lassa virus [20–22]. Thus, this robust Cas13a-based diagnostic tool may satisfy the requirements of *N. gonorrhoeae* detection and antibiotic resistant-SNPs identification in clinical practice.

Azithromycin combined with ceftriaxone is the currently recommended treatment for *N. gonorrhoeae* given the increasing MIC of ceftriaxone [23] and the emergence of high-level azithromycin resistance strains in various geographical regions showing a tendency to spread to other areas [24–30]. The main cause of elevated azithromycin resistance has been highly correlated with A2059G and C2611T mutations in the 23S rRNA subunit of the bacterial ribosome [31–33]. Herein, we attempt to develop and apply this Cas13a-based method to develop a rapid and accurate assay for *N. gonorrhoeae* detection and azithromycin resistance identification that will contribute to rational drug use by clinicians.

## Materials and Methods

### Ethics approval

All human samples used for this study were evaluated and approved by the Ethics Review Committee at Dermatology Hospital of Southern Medical University (2020056).

### Culture and azithromycin susceptibility testing

The *N. gonorrhoeae* strains used to verify azithromycin resistance were isolated from clinical samples of patients from Guangzhou, China and were identified by Gram Staining, oxidase, catalase, and sugar fermentation tests. Isolates were cultured in Thayer-Martin medium and incubated in 5% CO2 in a 37°C incubator.

Antimicrobial susceptibility to azithromycin was tested using the agar dilution method, according to WHO recommendations [34]. Briefly, all strains were cultured for 18 h and adjusted to a 0.5 McFarland standard suspension, and the cultures were then multipoint inoculated onto antimicrobial agar plates containing different drug concentrations.

### Sample preparation

Clinical urine samples were collected and stored at 4°C temporarily before DNA extraction. Urethral swabs were collected and stored in DNA/RNA Shield (Zymo Research; R1100-250) and 4°C for temporary storage before DNA extraction. Genome DNA extraction was carried out using the HiPure Bacterial DNA Kit (Magen; D3146-02) according to the manufacturer’s instructions. Extracted genomic DNA was stored at −20°C until use.

Clinical urine specimens for Roche COBAS 4800 NG/CT tests were prepared according to Roche’s standard operation protocols.

Serial dilutions of dsDNA (porA, A2059G, C2611T) were prepared by using Q5 High-Fidelity DNA Polymerases (New England Biolabs; M0492S) to amplify the target gene in a total reaction volume of 50 μL (25 μL of 2× Master Mix, 2.5 μL of each 10 μM primer, 2 μL of DNA template, and 18 μL ddH2O). PCR was performed as follows: initial denaturation at 98°C for 30 s, then 35 cycles at 98°C for 10 s, 60°C for 20 s, and 72°C for 30 s, followed by 72°C for 2 min. After amplification, PCR products were verified and purified following agarose gel electrophoresis. The Universal DNA purification kit (TIANGEN; DP214) was used according to the manufacturer’s protocol to extract target DNA from agarose gel. Purified dsDNA was quantified using the Qubit dsDNA HS Assay Kit (ThermoFisher; Q33230) and stored at −20°C until use. PCR primers used to produce dsDNA are reported in **Table S2**.

### Protein expression and purification of Cas13a

LwCas13a expression and purification were carried out according to the protocols described by Zhang et al. with some modifications [35]. Briefly, LwCas13a expression vectors (NovoPro Bioscience; V010159) were transformed into Rosetta (DE3) Competent Cells (Tiangen; CB108). Competent cells containing LwCas13a vectors were inoculated into LB Broth media (Sangon Biotech; A507002) containing 50 μg/mL ampicillin (Sangon Biotech; A100339) and grown at 37°C, 220 rpm until the OD600 reached 0.6. Isopropyl-beta-D-thiogalactopyranoside (Sangon Biotech; B541007) was added to the media at a final concentration of 0.5 mM to induce protein expression. Cells were then centrifuged at 4°C, and cell pellets were harvested and stored at −80°C for further purification.

Protein purification was performed at 4°C. Cell pellets were crushed and purified with His-tag Protein Purification Kit (Beyotime Biotechnology; P2226). Then incubated at 25°C for 3 h with SUMO protease (Novoprotein; PE007-01A) to digest SUMO tag. Purified Cas13a protein was stored at −80°C in storage buffer (50 mM Tris, 600 mM NaCl, 5% glycerol, 2 mM DTT, pH 7.5). All purification steps were analyzed and confirmed by SDS-PAGE and Coomassie Blue staining (Sangon Biotech; C510041). The concentration of protein was quantified using the BCA Protein Assay Kit (Beyotime Biotechnology; P0012S).

### crRNA preparation

For crRNA preparation, oligonucleotides containing the T7 promoter sequence, spacers (complement to target RNA), and the crRNA core sequence (bind to Cas13a) were designed by SnapGene 4.1.9 and NCBI BLAST, and synthesized by Sangon Biotech. Synthetic ssDNA (100 μM) binds to short T7 primer sequence (100 μM) by gradient annealing from 95°C to 25°C with cooling rate of 0.1°C/s. The product was then transcribed to crRNA using HiScribe T7 Quick High Yield RNA Synthesis kit (New England Biolabs; E2050S) incubating at 37°C overnight. Transcribed crRNA was purified using RNA XP Clean Beads (Beckman; A63987), and the concentration was quantified using the Qubit RNA HS Assay Kit (ThermoFisher; Q32852). crRNA was stored at −20°C until use. All crRNA used in this study are reported in **Table S2**.

### Recombinase Polymerase Amplification

Recombinase Polymerase Amplification (RPA) primers were designed by SnapGene 4.1.9 and NCBI BLAST according to the TwistAmp Assay Design Manual instructions, which can be downloaded from the official website (https://www.twistdx.co.uk/en/support/manuals/twistamp-manuals), with the condition that primers must flank the crRNA target region. RPA primers were synthesized by Sangon Biotech. The forward primer contained the T7 promoter sequence for initiating the transcription. TwistAmp Basic (TwistDx; TABAS03KIT) was used to amplify the target DNA. In a total reaction volume of 25 μL (containing 1.2 μL of each 10 μM primer, 14.75 μL of rehydration buffer, 1.25 μL of Magnesium Acetate (MgOAc), 1 μL of input, and 5.6 μL ddH2O). The mixture was run at 37°C for 2 h and then subjected to Cas13a detection assays. All RPA primers used in this study are available in **Table S2**.

### LwCas13a collateral detection

Cas13a detection assays were mainly performed according to the protocol described by Zhang et al. [19]. Briefly, the assay was carried out in a 25 μL reaction volume consisting of 40 mM Tris-HCl (pH7.5), 9 mM MgCl_2_, 1 mM rNTPs (New England Biolabs; N0466L), 50 U RNase inhibitor (New England Biolabs; M0314L), 37.5 U T7 RNA Polymerase (New England Biolabs; M0251L), 225 nM crRNA, 45nM purified LwCas13a, 125 nM RNA reporter (5′-6FAM-UUUUU-BHQ1-3′ as described by Gootenberg et al.[36]), and 1.25 μL RPA reaction solution was added to the above mixture. The reaction mixture was allowed to incubate at 37°C for 3 h in a 96-Well Half-Area Microplate (Corning; CLS3694-100EA). Fluorescence emission (excitation 490 nm/detection 520 nm) was tested every 5 min.

### Lateral flow readout

The lateral flow dipstick (Milenia Biotec; MGHD 1) was used to acquire the results of Cas13a collateral cleavage, which was based on the cleavage of the FITC-RNA-Biotin reporter. It basically replaces the RNA reporter used in the system described above, with a new RNA reporter (5′-FITC-UUUUUUUUUUUUUU-Biotin-3′ described by Myhrvold et al. [37]), and then was subjected to the same process. Subsequently, a 20 μL volume of Cas13a detection solution was added and the reaction mixture was incubated at 37°C for 3h in 80 μL dipstick buffer, with thorough mixing. A lateral flow dipstick was inserted into the mixture to obtain the results.

### Sanger sequencing

The 23s rRNA gene of *N. gonorrhoeae,* which contains the A2059G and C2611T point mutations, was amplified using the Q5 High-Fidelity DNA Polymerases (New England Biolabs; M0492S). In a 25 μL reaction volume, comprised of 12.5 μL of 2× Master Mix, 1.25 μL of each 10 μM primer, 1 μL of input, and 8 μL ddH2O. PCR was performed as follows: initial denaturation at 98°C for 30 s, then 35 cycles of 98°C for 10 s, 64°C for 20 s, and 72°C for 30 s, followed by 72°C for 2 min. PCR products were verified by Sanger sequencing (Sangon Biotech). The results of sequencing were blasted in SnapGene 4.1.9. PCR primers are available in **Table S2**.

### Analysis of fluorescence data

Prism 8 software (GraphPad, La Jolla, CA, USA) was used for visualization of results and data analyses. Data are presented as mean ± SEM and were tested for normality with the Shapiro-Wilk test. Differences were considered significant at *P-*values < 0.05.

## Results

### Schematic of Cas13a based *N. gonorrhoeae* detection and azithromycin resistance identification

The SHERLOCK assay was performed as established by Zhang et al. [19], and combined RPA and Cas13a to create an isothermal detection system. The target sequence was amplified by RPA, and the T7 promoter was appended to the front of the RPA product as the template to be used to initiate subsequent RNA transcription. Synthetic crRNA guided the Cas13a protein to recognize the specific target and enable its RNA cleavage and collateral cleavage activities (**Fig. 1**) [18]. For *N. gonorrhoeae* detection, we selected porA as the target as was frequently used to identity *N. gonorrhoeae* in other methods [17, 38–40]. Based on the characteristics of its single-base resolution, we constructed two crRNA sequences that could identify A2059G and C2611T separately. Although a single synthetic mismatch of crRNA pairing to the target enabled the assay to identify the A2059G mutation (**Fig. S1A**), this design failed to achieve the identical results for the detection of the C2611T mutation. Thus, we introduced one more synthetic mismatch of crRNA for C2611T mutation testing (**Fig. S1B-D**). Both crRNA designs were successfully utilized for 23s rRNA mutations detection.

**Fig. 1.**
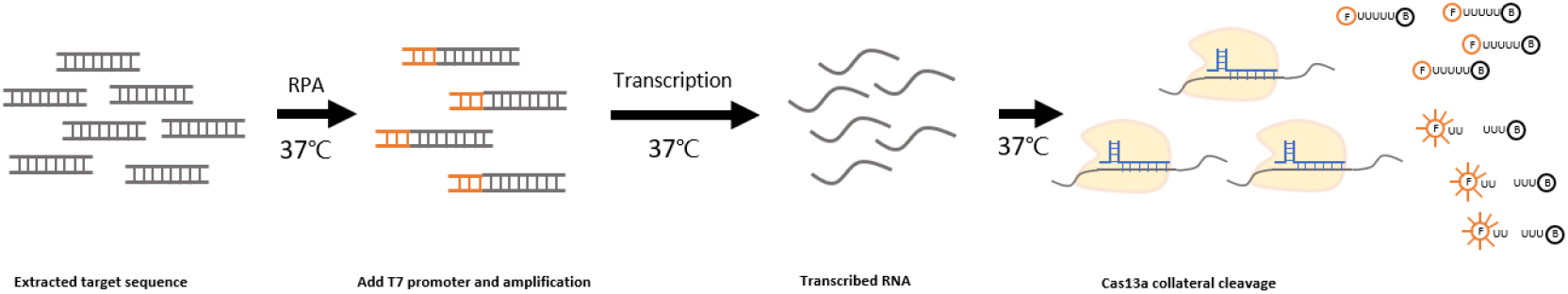
Schematic overview of the SHERLOCK assay. Illustration of LwCas13a combined with RPA for detection of *N. gonorrhoeae.* The target gene was amplified by RPA and appended with the T7 promoter. The unpurified RPA product is added to the Cas13a system. Once crRNA was matched the transcribed RNA target, Cas13a cleaves the RNA reporter and the reporters emit a fluorescent signal.

### Evaluation of the Cas13a-based method with sensitivity and specificity

To determine the sensitivity of the *N. gonorrhoeae* detection and azithromycin resistance identification assay, we prepared serial dilutions of dsDNA template, ranging from 10^0^ copy/μL to 10^5^ copies/μL (**Table 1**). The RPA step included the addition of 1 μL input of dsDNA template, which was then transferred to the mixture for Cas13a detection. Detection of 10 copies/μL was achieved for porA and C2611T identification (**Fig. S2A, S2E**). For A2059G identification, the detection limit was an order of magnitude lower, at about 10^2^ copies/μL (**Fig. S2C**). We further evaluated the specificity of the assay using a panel of urogenital tract pathogenic bacteria (n=12) and *Neisseria meningitides* (**Table 2**). No cross-reactivity was observed for both *N. gonorrhoeae* detection and azithromycin resistance identification (**Fig. S2B, S2D, S2F**). All RPA primers and crRNA sequences had been confirmed by BLAST before we tested its specificity. The SHERLOCK exhibits robust capability for *N. gonorrhoeae* detection and azithromycin resistance identification.

**Table 1.** LODs of porA detection and 23S rRNA point mutation identification (copies per reaction)

| Assay | 10<sup>4</sup> | 10<sup>3</sup> | 10<sup>2</sup> | 10<sup>1</sup> | 10<sup>0</sup> | Wild type<sup>a</sup> |
| --- | --- | --- | --- | --- | --- | --- |
| porA | 3/3 | 3/3 | 3/3 | 3/3 | 0/3 | N/A |
| A2059G | 3/3 | 3/3 | 3/3 | 0/3 | 0/3 | 0/3 |
| C2611T | 3/3 | 3/3 | 3/3 | 3/3 | 0/3 | 0/3 |

<sup>a</sup>N/A, not applicable

**Table 2.** Specificity of porA detection and 23S rRNA point mutation identification

| Organism | Assay<sup>a</sup> |  |  |
| --- | --- | --- | --- |
|  | porA | A2059G | C2611T |
| porA | Pos | N/A | N/A |
| A2059G template | N/A | Pos | Neg |
| C2611T template | N/A | Neg | Pos |
| *Neisseria Meningitidis* | Neg | Neg | Neg |
| *Treponema pallidum* | Neg | Neg | Neg |
| *Herpes Simplex Virus-1* | Neg | Neg | Neg |
| *Herpes Simplex Virus-2* | Neg | Neg | Neg |
| *Human Papilloma Virus-2* | Neg | Neg | Neg |
| *Human Papilloma Virus-7* | Neg | Neg | Neg |
| *Candida Albicans* | Neg | Neg | Neg |
| *Trichomonas Vaginalis* | Neg | Neg | Neg |
| *Chlamydia Trachomatis* | Neg | Neg | Neg |
| *Ureaplasma Urealyticum* | Neg | Neg | Neg |
| *Mycoplasma Humanum* | Neg | Neg | Neg |
| *Escherichia Coli* | Neg | Neg | Neg |
| *Enterococcus Faecalis* | Neg | Neg | Neg |

<sup>a</sup>Pos, positive; Neg, negative; N/A, not applicable.

### Validation of *N. gonorrhoeae* detection in clinical urine samples

Twenty-three clinical urine samples with low concentrations previously tested using the standard procedure of the Roche Cobas 4800 NG/CT test were used to validate the performance of SHERLOCK. DNA was extracted from urine samples after centrifugation and a maximum volume of 6.6 μL of DNA was amplified by RPA, followed by Cas13a detection. The method was repeated using 3 technical replicates and the fluorescence signal of each sample was normalized against the negative controls. Using this method, a total of 12 of 23 positive samples were detected, showing a 100% coincidence rate with the Roche assay (**Fig. 2**). The fluorescence signals of 3 samples (samples 3, 6, 10) were weaker than other specimens, but still could be distinguished with the negative control.

**Fig. 2.**
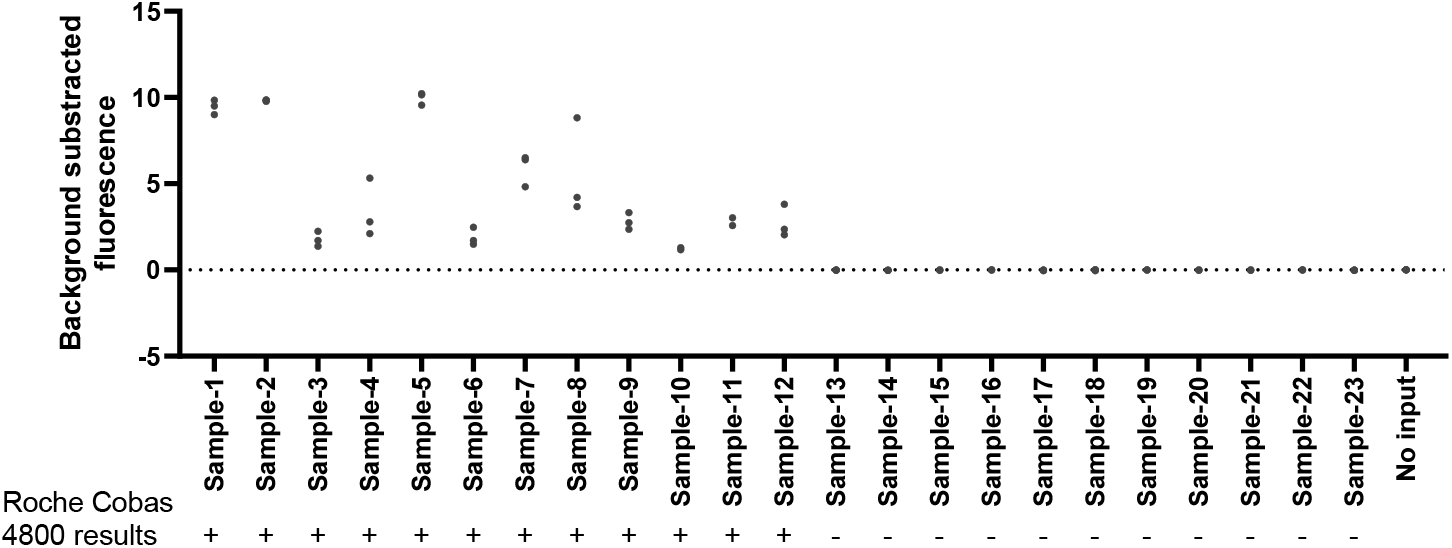
Validation of the SHERLOCK assay for urine samples. The performance of SHERLOCK assay in clinical urine samples compared to verification using the Roche Cobas 4800 (NG/CT test).

### Validation of azithromycin resistance identification

A2059G mutant strains (n=8), C2611T mutant strains (n=8), and wild-type strains (n=8) isolated from clinical specimens were used to validate Cas13a-based SNPs detection. We measured the MICs of azithromycin in each strain and sequenced their 23S rRNA gene (**Fig. 3, 4**). Strains containing either the A2059G or C2611T point mutation were more likely to be a high-level azithromycin-resistant strain. Of the 8 A2059G isolated strains, all 8 strains had MICs of >1 mg/L, and 5 A2059G strains had MICs ≥512 mg/L. Of the 8 C2611T isolated strains, 7 had MICs of >1 mg/L and 3 C2611T strains had MICs ≥512 mg/L. Compared with mutant strains, the wild-type strains possessed lower MICs, corresponding to ≤1 mg/L in 7 strains and the MIC of the remaining strain was 4 mg/L, which was above the average MICs of all mutant strains. We extracted DNA of all mutant and wild-type strains to validate this Cas13a-based assay. Paired with A2059G crRNA or C2611T crRNA, this assay successfully differentiated all 8 A2059G mutant strains and 8 C2611T strains from 8 wild-type strains (**Fig. 3, 4**). Sixteen strains harboring A2059G and C2611T mutations were identified by exhibiting a higher fluorescence intensity than wild-type strains by our assay (**Fig. S3A, S3B**). Thus, we successfully applied Cas13a based assay in azithromycin resistance identification.

**Fig. 3.**
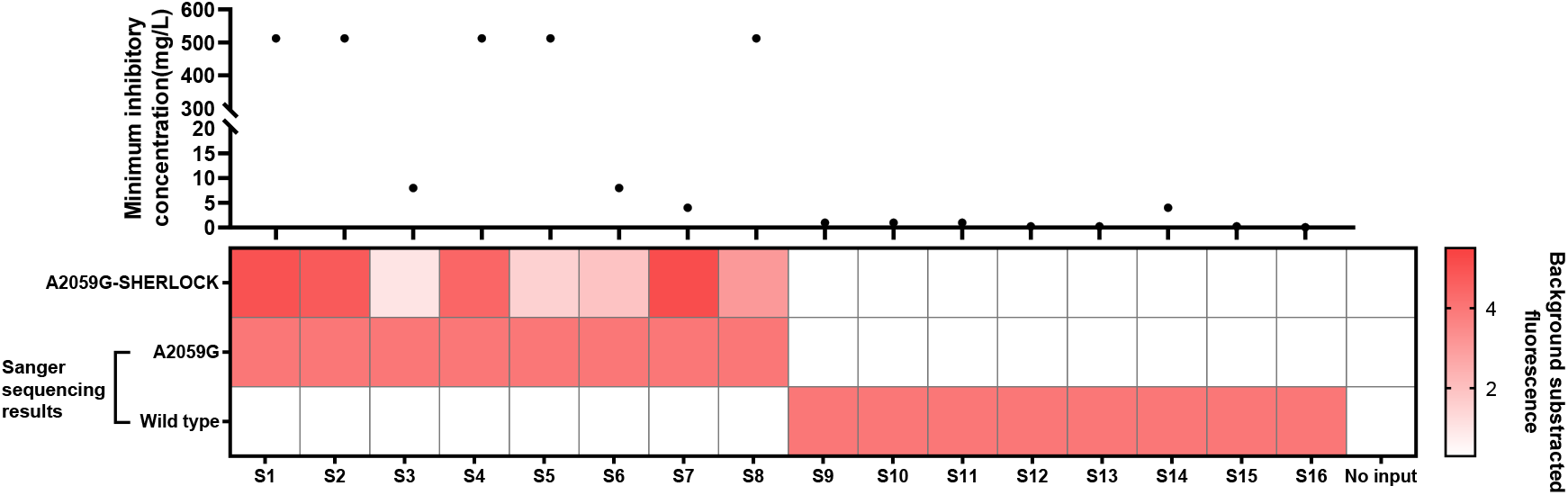
Identification of the SHERLOCK assay for the identification of the A2059G point mutation. Sixteen *N. gonorrhoeae* strains were tested to measure their MICs for azithromycin and then assayed for 23S rRNA for 2059 and 2611 point mutations. Extracted DNA was tested by SHERLOCK assay directly.

**Fig. 4.**
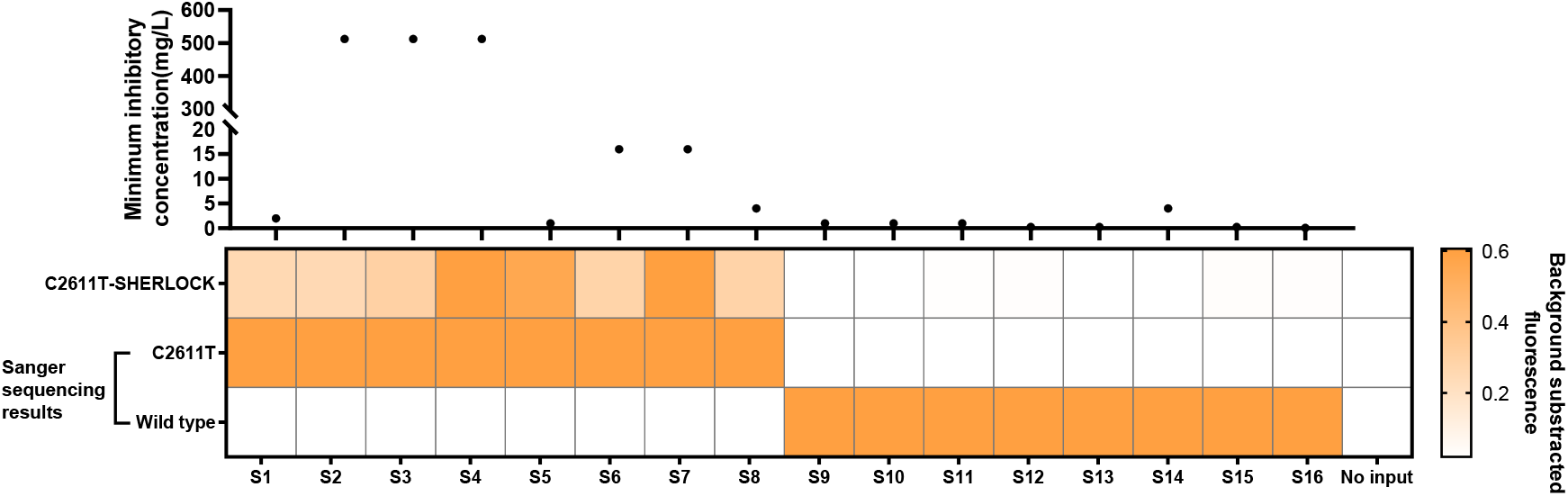
Evaluation of the SHERLOCK assay for C2611T point mutation detection. Sixteen *N. gonorrhoeae* strains were evaluated to determine MICs of azithromycin and were then assayed for 23S rRNA 2059 and 2611 point mutations. Extracted DNA was tested by SHERLOCK assay directly.

### Lateral flow for *N. gonorrhoeae* detection and azithromycin resistance identification

We applied lateral flow to provide a more convenient readout tool. The FAM and BHQ1 markers in the RNA reporter were replaced by FITC and Biotin. Compared to the fluorescent intensity detection, the lateral flow is inserted directly into the reaction liquid instead of using a specific device or instrument for the readout of results. The lateral flow contains a control band and a test band. Generally, a positive test will show only one test band or two bands (test band and control band), due to its different cleavage efficiency which will result in varying amounts of cleaved RNA reporter captured by the antibody in the test band. We tested lateral flow for porA detection, A2059G identification, and C2611T identification separately (**Fig. 5A-C**). Three positive groups generated visual signals in the test bands, while all wild-type groups and the no-input group only showed a single control band.

**Fig. 5.**
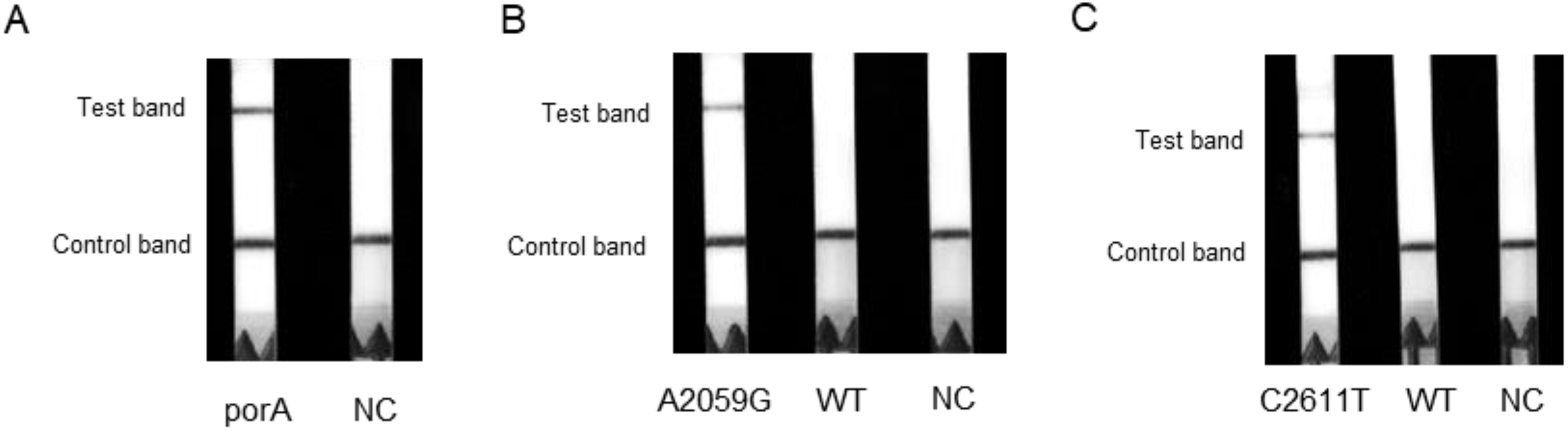
The SHERLOCK assay using lateral flow readout for the detection of *N. gonorrhoeae*. (A) Lateral flow readout system applied to the SHERLOCK assay for the detection of the porA gene. (B) Lateral flow result for the detection of the A2059G point mutation. (C) Lateral flow result for the detection of the C2611T point mutation.

### Applying Cas13a based *N. gonorrhoeae* detection and azithromycin resistance identification in urethritis

To confirm the efficacy of the assay in clinical specimens. We collected 27 urethral swabs from patients with urethritis requiring differential diagnosis for potential gonococcal infection and to determine whether azithromycin was still effective and this information is important for the clinician. We extracted DNA from urethral swabs directly and then tested all 27 samples with the SHERLOCK assay (**Fig. 6**). Overall, 62.96% (17/27) samples showed porA positivity, and the fluorescence intensity of 17 samples was higher than that of the negative samples and the no-input group. We further tested for azithromycin resistance with A2059G crRNA and C2611T crRNA, and no mutant strain was discovered in the 17 porA positive samples. For 27 specimens, we sequenced the 23S rRNA gene and the results showed a 100% coincidence rate with our assay (**Table S1**). Sequencing data demonstrated that 17 porA positive samples were wild-type strains and no signals were detected in 10 porA negative samples.

**Fig. 6.**
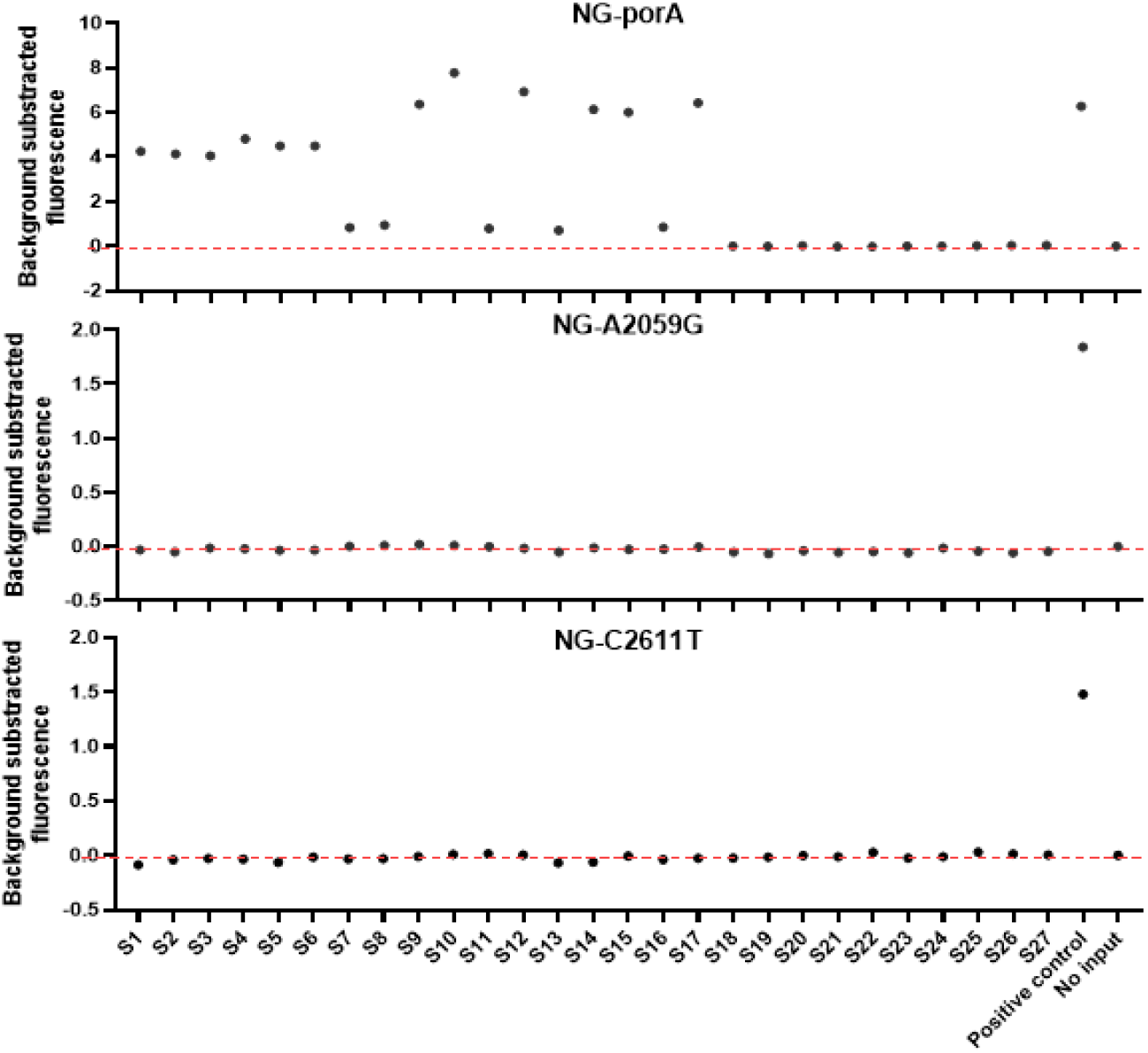
Fluorescence intensity of *N. gonorrhoeae* SHERLOCK assay for the detection of gonococcal urethritis. The performance of the CRISPR/Cas13a-based assay for using urethral swab samples from patients with urethritis for identification of porA, A2059G, and C2611T.

## Discussion

Antimicrobial resistance is the largest concern in the treatment of gonorrhea, with its decreasing susceptibility to antibiotics used in previous or current treatment approaches [5]. Treatment failures have been reported to the current first-line drug combination of azithromycin and ceftriaxone [8, 9], and represent a significant treatment challenge to clinicians. Rapid information regarding antimicrobial resistance would be beneficial to rational drug use in the clinic and would slow this growing trend. Traditional culture-based drug resistance methods have been widely used in the clinical laboratory and are of great importance in antimicrobial resistance surveillance, although the testing is time-consuming and is unable to meet clinical requirements rapidly [41]. Newly NAATs-based assays may overcome some of the disadvantages of culture-based methods and can be applied to identify antimicrobial resistance associated mutations simultaneously. Current molecular methods to identify mutations are mainly based on specially designed probes, HRM analysis, and mass spectrometry [13, 16, 17]. Compared with culture-based antimicrobial resistance detection methods, these methods effectively shorten assay times, but still present several limitations. If the alteration in gene has little impact on Tm values or GC content, for example, C to G variations, the method would be invalidated, moreover, short insertions or deletions may make the method unreliable [17, 42]. Furthermore, these assays are essentially PCR-based and require PCR amplification instruments coupled with other detection equipment, which limits the convenience and flexibility of the assay. Thus, the development of an assay with ultra-high resolution is desired for distinguishing mutant from the wild-type.

CRISPR/Cas molecular diagnostics have been developed and applied for testing various organisms, including SARS-CoV-2, HPV, Zika virus, Dengue virus, Ebola virus, and plasmodium [20, 21, 37, 43, 44]. Benefiting from its high specificity, sensitivity, and ability to identify SNP with the isothermal process, we have adopted a Cas13a-based strategy for *N. gonorrhoeae* detection and antimicrobial resistance identification in this study. The SHERLOCK contains two nucleic acid amplification steps: DNA amplification by recombinase polymerase amplification and RNA amplification by T7 transcription. With double signal amplification cycles, this strategy allowed to detect low levels of *N. gonorrhoeae*. Cross-reactivity is the major concern of currently developed diagnostic methods [12]. Attempts have been made to introduce two targets to uniquely identify a species, but this approach complicates the assay [13, 15–17]. The combination of specially-designed RPA primers and crRNA makes the whole reaction extremely specific. As expected, our assay exhibited high specificity in testing a panel of non-gonococcal bacteria. In addition, our Cas13a-based assay showed an excellent concordance rate with the Roche Cobas assay currently used for clinical urine samples. With regard to SNPs detection, assay has been developed that exploits CRISPR/Cas13a technology to recognize single point mutations [19, 36, 45]. Our Cas13a-based assay achieved a sensitivity of 10 copies per reaction, which is more sensitive than previous HRM-based assays [46]. The diagnostic capability of the Cas13a-based assay has also been examined by testing clinical isolates harboring the SNP mutation. This isothermal assay which relies on a reaction temperature of 37°C over the entire process without complex equipment has a great potential to be applied as a POCT device.

Azithromycin is a widely used macrolide antimicrobial agent and primarily acts on domain V of the 23S rRNA gene. Previous studies and our observation have revealed that *N. gonorrhoeae* strains harboring A2059G and C2611T mutations in the 23s rRNA gene are strongly associated with high-level azithromycin resistance [28, 32, 33, 47–49]. Sixteen *N. gonorrhoeae* isolates containing 23s rRNA mutations were utilized to evaluate the performance of our assay. The results showed that our Cas13a-based assay could provide drug resistance information in real-time. We also tested a small number of urethral swabs collected from the clinic in Guangzhou, China. In porA-positive(17/27) urethral swabs from patients with urethritis, no 23s rRNA mutant was identified, which is consistent with previous reports, indicating that the high-level azithromycin-resistant *N. gonorrhoeae* has not wildly spread in Guangzhou, China [14, 16, 17]. Because of the ongoing use of ceftriaxone and azithromycin dual therapy, the surveillance of 23s rRNA mutation is still a requirement.

There are several limitations to our study. We only tested a small number of samples and limited sources of clinical specimen. Cervical, anus, and pharynx specimens should be considered in further study. Moreover, the cost of SHERLOCK is higher than HRM-based method, though lower than most reported assays to date. In summary, we developed a CRISPR/Cas13a-based assay for *N. gonorrhoeae* detection and azithromycin resistance identification with great potential for providing drug resistance information to assist clinical diagnosis and treatment.

## Acknowledgments

This work was supported by grants from the Overseas Famous Teacher Project of Guangdong Provincial Department of Science and Technology (No. 2020A1414010136), Medical Science and Technology Research Fundation of Guangdong Province (No. A2019010 and No. A2021139), Guangdong Traditional Chinese Medicine Research Project (No. 20191230 and No. 20211277), Guangdong Provincial Medical Research Fund (No. B2020149), Scientific Research Initiative Project of Southern Medical University (Project of Youth Science and Technology Personnel Training, No. PY2018N100), Key scientific research platforms and research projects of colleges and universities in Guangdong Province (No. 2018KQNCX025). The funders had no role in study design, data collection and analysis, decision to publish, or preparation of the manuscript.

## Author contributions

Conceived and designed the study: HL and HPZ. Collected samples: ZDM, XML, JJY, JLO, QQX, and ZQF. Isolation of clinical strains: LHZ, YYP, QHX. Tested Susceptibility to azithromycin: XML, XXL, and YWL. Performed the laboratory work: HL. Analyzed the data: HL and WC. Wrote the initial draft of the paper: HL and WC. Funding supported the study: XLQ, XML, and HPZ. All authors viewed and contributed to the final paper.

---

**Fig. S1.**
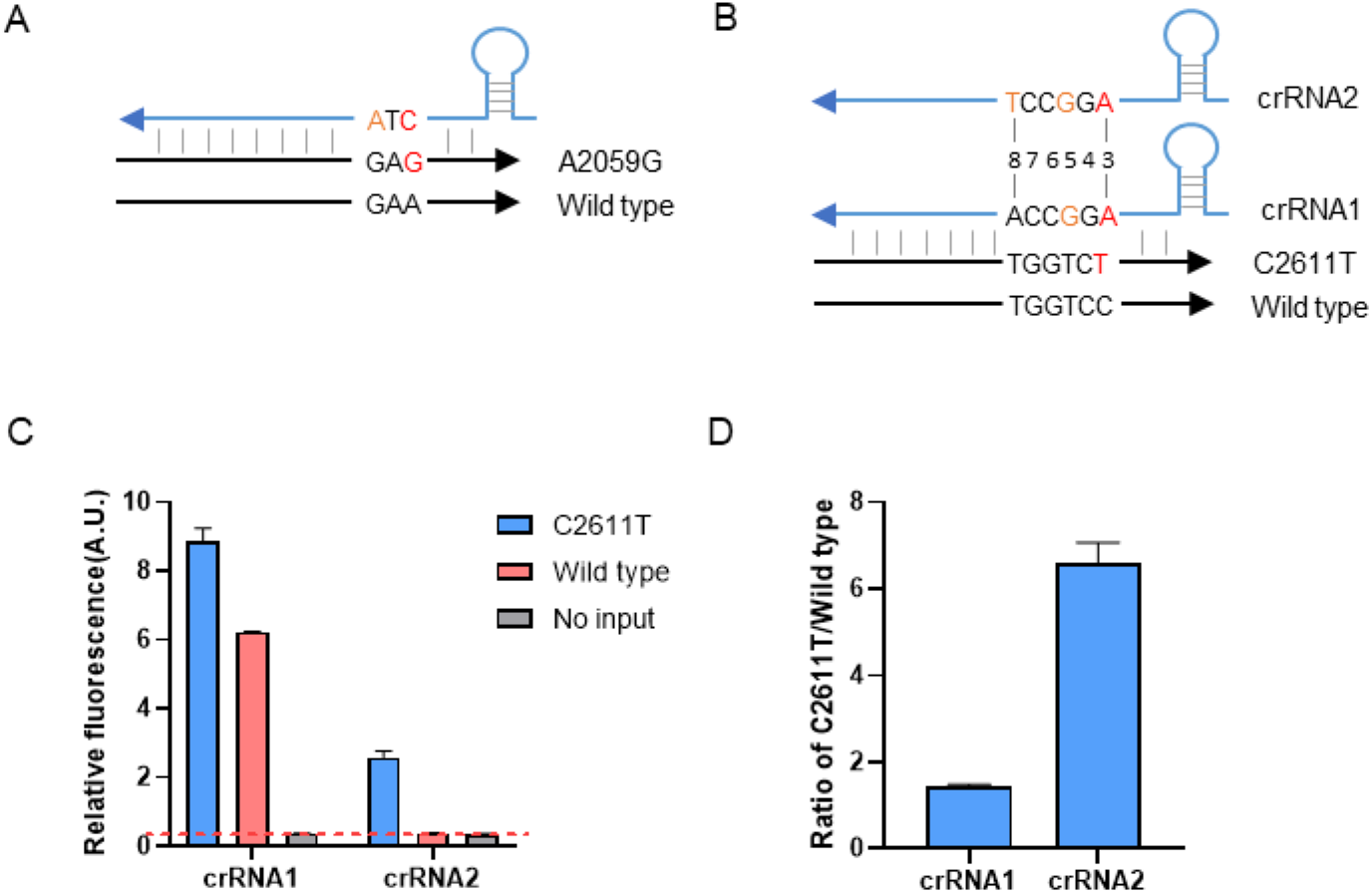
Schematic diagram illustrating the crRNA design for the detection A2059G and C2611T. (A) crRNA for the identification of A2059G and one synthetic mismatches are highlighted. (B) crRNA for the identification of C2611T; one synthetic mismatches are highlighted in crRNA1 and two synthetic mismatches are highlighted in crRNA2. (C) Fluorescence intensity of crRNA1 and crRNA2 for the detection of C2611T and wild-type template (n=3 technical replicates). (D) The ratio of fluorescence intensity of C2611T to WT.

**Fig. S2.**
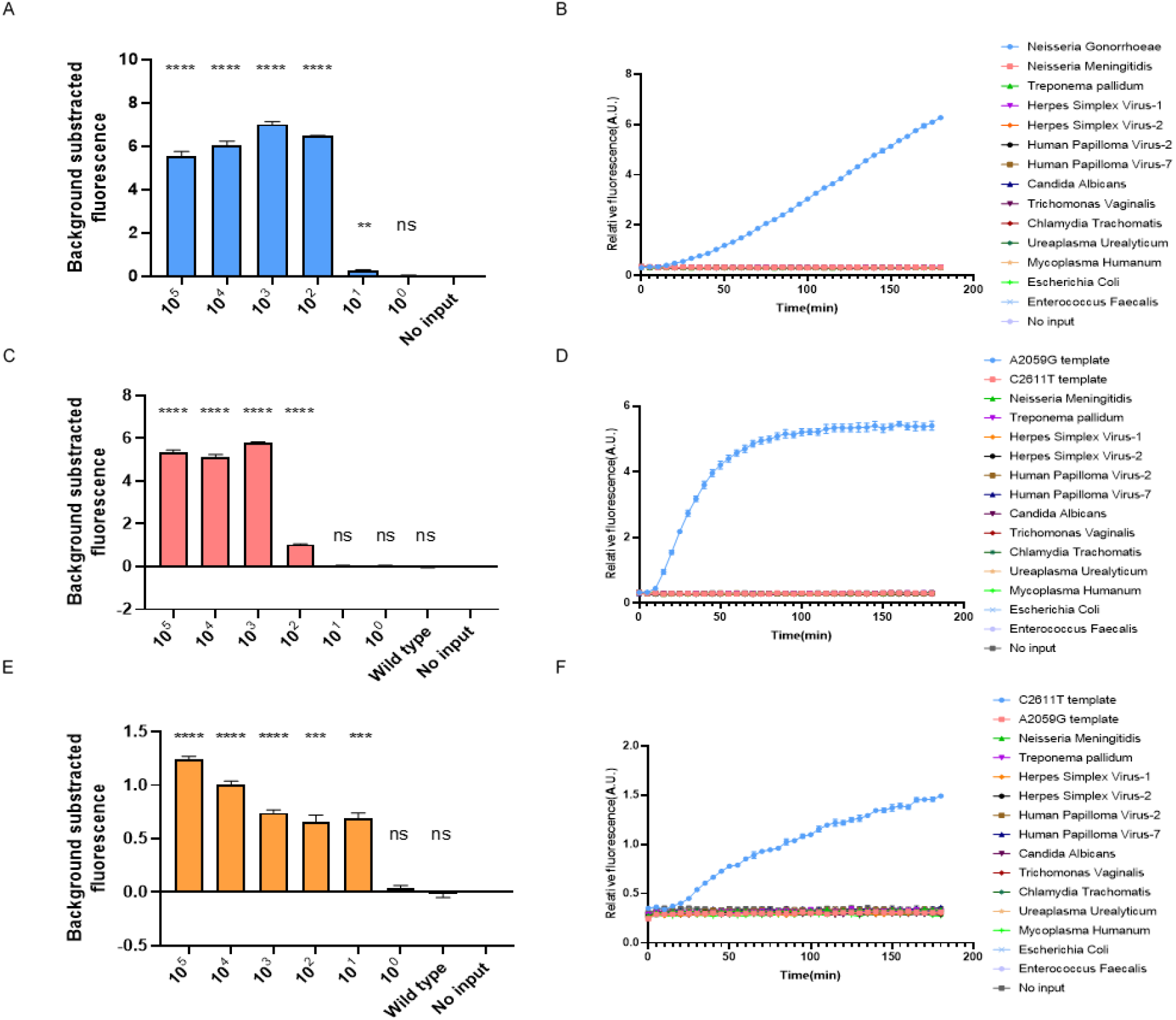
Sensitivity and specificity of the SHERLOCK assay for the detection of porA, A2059G, and C2611T. (A) Sensitivity of porA gene detection (n=3 technical replicates, two-tailed Student t-test; ns, not significant; **, p < 0.01; ****, p < 0.0001; bars represent mean ± SEM). (B) Specificity of porA detection (n=3 technical replicates). (C) Sensitivity of A2059G detection (n=3 technical replicates, two-tailed Student t-test; ns, not significant; ****, p < 0.0001; bars represent mean ± SEM). (D) Specificity of A2059G detection (n=3 technical replicates). (E) Sensitivity of C2611T detection (n=3 technical replicates, two-tailed Student t-test; ns, not significant; ***, p < 0.001; ****, p < 0.0001; bars represent mean ± SEM). (F) Specificity of C2611T detection (n=3 technical replicates).

**Fig. S3.**
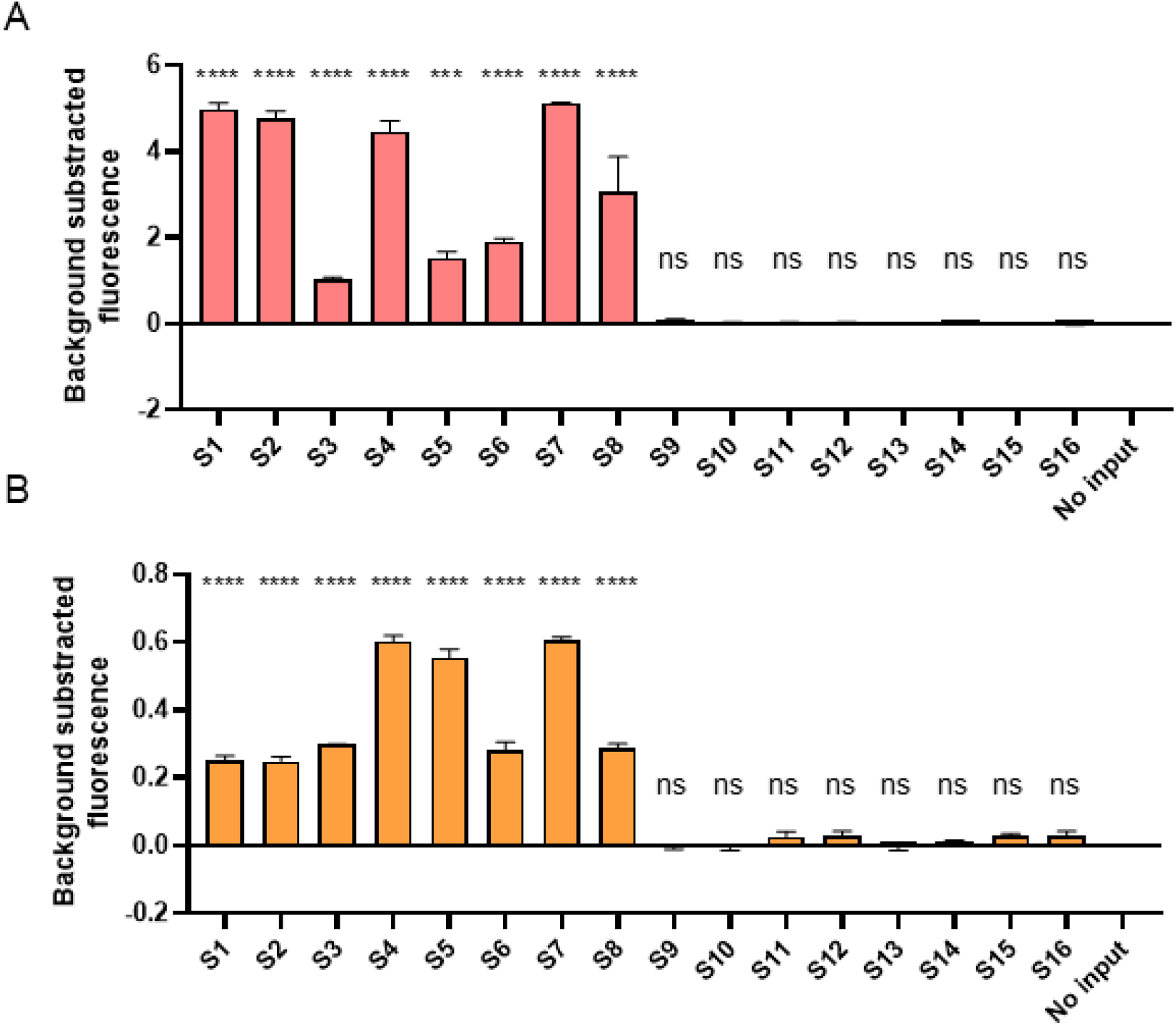
Fluorescence intensity relative to azithromycin resistance mutation strain detection. (A) Fluorescence intensity of the SHERLOCK assay for the detection of 16 *Neisseria gonorrhoeae* strains (8 A2059G mutant strains and 8 wild-type strains) for A2059G detection. (n=3 technical replicates, two-tailed Student t-test; ns, not significant; *, p < 0.05, ***, p < 0.001; ****, p < 0.0001; bars represent mean ± SEM). (B) Fluorescence intensity of 16 *N. gonorrhoeae* strains (8 C2611T mutant strains and 8 wild-type strains) for C2611T detection. (n=3 technical replicates, two-tailed Student t-test; ns, not significant; ****, p < 0.0001; bars represent mean ± SEM)

**Table S1.** Applying Neisseria gonorrhoeae SHERLOCK for detection of urethritis

| Sample ID | Clinical diagnosis | Assay<sup>a</sup> |  |  | Sanger sequencing<sup>a,b</sup> |
| --- | --- | --- | --- | --- | --- |
|  |  | poA | A2059G | C2611T |  |
| S1 | urethritis | Pos | Neg | Neg | WT |
| S2 | urethritis | Pos | Neg | Neg | WT |
| S3 | urethritis | Pos | Neg | Neg | WT |
| S4 | urethritis | Pos | Neg | Neg | WT |
| S5 | urethritis | Pos | Neg | Neg | WT |
| S6 | urethritis | Pos | Neg | Neg | WT |
| S7 | urethritis | Pos | Neg | Neg | WT |
| S8 | urethritis | Pos | Neg | Neg | WT |
| S9 | urethritis | Pos | Neg | Neg | WT |
| S10 | urethritis | Pos | Neg | Neg | WT |
| S11 | urethritis | Pos | Neg | Neg | WT |
| S12 | urethritis | Pos | Neg | Neg | WT |
| S13 | urethritis | Pos | Neg | Neg | WT |
| S14 | urethritis | Pos | Neg | Neg | WT |
| S15 | urethritis | Pos | Neg | Neg | WT |
| S16 | urethritis | Pos | Neg | Neg | WT |
| S17 | urethritis | Pos | Neg | Neg | WT |
| S18 | urethritis | Neg | Neg | Neg | N/A |
| S19 | urethritis | Neg | Neg | Neg | N/A |
| S20 | urethritis | Neg | Neg | Neg | N/A |
| S21 | urethritis | Neg | Neg | Neg | N/A |
| S22 | urethritis | Neg | Neg | Neg | N/A |
| S23 | urethritis | Neg | Neg | Neg | N/A |
| S24 | urethritis | Neg | Neg | Neg | N/A |
| S25 | urethritis | Neg | Neg | Neg | N/A |
| S26 | urethritis | Neg | Neg | Neg | N/A |
| S27 | urethritis | Neg | Neg | Neg | N/A |

<sup>a</sup>Pos, positive; Neg, negative; N/A, not applicable.
<sup>b</sup>WT, 23S rRNA wild type strain

**Table S2.**
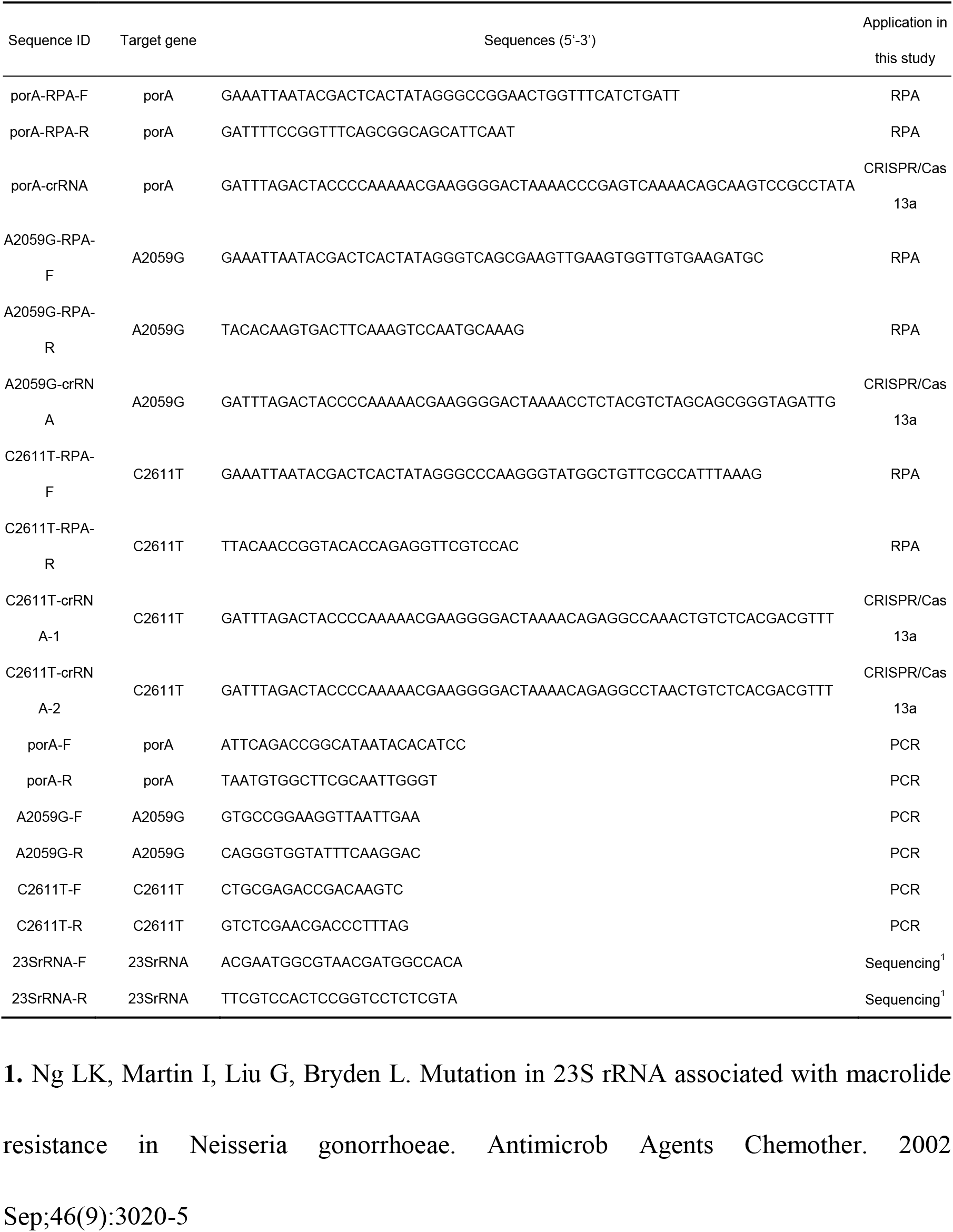
Primer and crRNA sequences

## Reference

1. Unemo M, Seifert HS, Hook EW, 3rd, Hawkes S, Ndowa F, Dillon JR. Gonorrhoea. Nat Rev Dis Primers 2019; 5(1): 79.

2. Rowley J, Vander Hoorn S, Korenromp E, et al. Chlamydia, gonorrhoea, trichomoniasis and syphilis: global prevalence and incidence estimates, 2016. Bull World Health Organ 2019; 97(8): 548–62p.

3. Newman L, Rowley J, Vander Hoorn S, et al. Global Estimates of the Prevalence and Incidence of Four Curable Sexually Transmitted Infections in 2012 Based on Systematic Review and Global Reporting. PLoS One 2015; 10(12): e0143304.

4. Rice PA, Shafer WM, Ram S, Jerse AE. Neisseria gonorrhoeae: Drug Resistance, Mouse Models, and Vaccine Development. Annu Rev Microbiol 2017; 71: 665–86.

5. Suay-García B, Pérez-Gracia MT. Future Prospects for Neisseria gonorrhoeae Treatment. Antibiotics (Basel) 2018; 7(2).

6. Unemo M, Shafer WM. Antimicrobial resistance in Neisseria gonorrhoeae in the 21st century: past, evolution, and future. Clin Microbiol Rev 2014; 27(3): 587–613.

7. WHO Guidelines Approved by the Guidelines Review Committee. WHO Guidelines for the Treatment of Neisseria gonorrhoeae. Geneva: World Health Organization Copyright World Health Organization 2016., 2016.

8. Eyre DW, Sanderson ND, Lord E, et al. Gonorrhoea treatment failure caused by a Neisseria gonorrhoeae strain with combined ceftriaxone and high-level azithromycin resistance, England, February 2018. Euro Surveill 2018; 23(27).

9. Whiley DM, Jennison A, Pearson J, Lahra MM. Genetic characterisation of Neisseria gonorrhoeae resistant to both ceftriaxone and azithromycin. Lancet Infect Dis 2018; 18(7): 717–8.

10. Gonococcal antimicrobial susceptibility surveillance in Europe, 2018. Available at: https://www.ecdc.europa.eu/en/publications-data/gonococcal-antimicrobial-susceptibility-surveillance-europe-2018. Accessed April.

11. Sexually Transmitted Disease Surveillance 2018. Available at: https://www.cdc.gov/std/stats18/gisp2018/default.htm. Accessed April.

12. Donà V, Low N, Golparian D, Unemo M. Recent advances in the development and use of molecular tests to predict antimicrobial resistance in Neisseria gonorrhoeae. Expert Rev Mol Diagn 2017; 17(9): 845–59.

13. Li Y, Xiu L, Liu J, et al. A multiplex assay for characterization of antimicrobial resistance in Neisseria gonorrhoeae using multi-PCR coupled with mass spectrometry. J Antimicrob Chemother 2020; 75(10): 2817–25.

14. Trembizki E, Buckley C, Donovan B, et al. Direct real-time PCR-based detection of Neisseria gonorrhoeae 23S rRNA mutations associated with azithromycin resistance. J Antimicrob Chemother 2015; 70(12): 3244–9.

15. Donà V, Smid JH, Kasraian S, et al. Mismatch Amplification Mutation Assay-Based Real-Time PCR for Rapid Detection of Neisseria gonorrhoeae and Antimicrobial Resistance Determinants in Clinical Specimens. J Clin Microbiol 2018; 56(9).

16. Peterson SW, Martin I, Demczuk W, et al. Multiplex real-time PCR assays for the prediction of cephalosporin, ciprofloxacin and azithromycin antimicrobial susceptibility of positive Neisseria gonorrhoeae nucleic acid amplification test samples. J Antimicrob Chemother 2020; 75(12): 3485–90.

17. Xiu L, Li Y, Wang F, et al. Multiplex High-Resolution Melting Assay for Simultaneous Identification of Molecular Markers Associated with Extended-Spectrum Cephalosporins and Azithromycin Resistance in Neisseria gonorrhoeae. J Mol Diagn 2020.

18. Abudayyeh OO, Gootenberg JS, Konermann S, et al. C2c2 is a single-component programmable RNA-guided RNA-targeting CRISPR effector. Science 2016; 353(6299): aaf5573.

19. Gootenberg JS, Abudayyeh OO, Lee JW, et al. Nucleic acid detection with CRISPR-Cas13a/C2c2. Science 2017; 356(6336): 438–42.

20. Lee RA, Puig H, Nguyen PQ, et al. Ultrasensitive CRISPR-based diagnostic for field-applicable detection of Plasmodium species in symptomatic and asymptomatic malaria. Proc Natl Acad Sci U S A 2020; 117(41): 25722–31.

21. Barnes KG, Lachenauer AE, Nitido A, et al. Deployable CRISPR-Cas13a diagnostic tools to detect and report Ebola and Lassa virus cases in real-time. Nat Commun 2020; 11(1): 4131.

22. Joung J, Ladha A, Saito M, et al. Detection of SARS-CoV-2 with SHERLOCK One-Pot Testing. N Engl J Med 2020; 383(15): 1492–4.

23. Update to CDC’s Sexually transmitted diseases treatment guidelines, 2010: oral cephalosporins no longer a recommended treatment for gonococcal infections. MMWR Morb Mortal Wkly Rep 2012; 61(31): 590–4.

24. Salmerón P, Moreno-Mingorance A, Trejo J, et al. Emergence and dissemination of three mild outbreaks of Neisseria gonorrhoeae with high-level resistance to azithromycin in Barcelona, 2016-18. J Antimicrob Chemother 2020.

25. Holderman JL, Thomas JC, Schlanger K, et al. Sustained Transmission of Neisseria gonorrhoeae with High-Level Resistance to Azithromycin, Indianapolis, Indiana 2017-2018. Clin Infect Dis 2021.

26. Shimuta K, Lee K, Yasuda M, et al. Characterization of two Neisseria gonorrhoeae strains with high-level azithromycin resistance isolated in 2015 and 2018 in Japan. Sex Transm Dis 2020.

27. Palavecino EL, Kilic A, Schmerer MW, Dobre-Buonya O, Toler C, McNeil CJ. First Case of High-Level Azithromycin-Resistant Neisseria gonorrhoeae in North Carolina. Sex Transm Dis 2020; 47(5): 326–8.

28. Liu YH, Wang YH, Liao CH, Hsueh PR. Emergence and Spread of Neisseria gonorrhoeae Strains with High-Level Resistance to Azithromycin in Taiwan from 2001 to 2018. Antimicrob Agents Chemother 2019; 63(9).

29. Gernert KM, Seby S, Schmerer MW, et al. Azithromycin susceptibility of Neisseria gonorrhoeae in the USA in 2017: a genomic analysis of surveillance data. Lancet Microbe 2020; 1(4): e154–e64.

30. Banhart S, Selb R, Oehlmann S, et al. The mosaic mtr locus as major genetic determinant of azithromycin resistance of Neisseria gonorrhoeae, Germany, 2018. J Infect Dis 2021.

31. Ng LK, Martin I, Liu G, Bryden L. Mutation in 23S rRNA associated with macrolide resistance in Neisseria gonorrhoeae. Antimicrob Agents Chemother 2002; 46(9): 3020–5.

32. Chisholm SA, Dave J, Ison CA. High-level azithromycin resistance occurs in Neisseria gonorrhoeae as a result of a single point mutation in the 23S rRNA genes. Antimicrob Agents Chemother 2010; 54(9): 3812–6.

33. Laumen JGE, Manoharan-Basil SS, Verhoeven E, et al. Molecular pathways to high-level azithromycin resistance in Neisseria gonorrhoeae. J Antimicrob Chemother 2021.

34. Organization WH. Manual for the laboratory identification and antimicrobial susceptibility testing of bacterial pathogens of public health concern in the developing word. Available at: http://www.who.int/csr/resources/publications/drugresist/en/IIAMRmanual.pdf?ua=1.

35. Kellner MJ, Koob JG, Gootenberg JS, Abudayyeh OO, Zhang F. SHERLOCK: nucleic acid detection with CRISPR nucleases. Nat Protoc 2019; 14(10): 2986–3012.

36. Gootenberg JS, Abudayyeh OO, Kellner MJ, Joung J, Collins JJ, Zhang F. Multiplexed and portable nucleic acid detection platform with Cas13, Cas12a, and Csm6. Science 2018; 360(6387): 439–44.

37. Myhrvold C, Freije CA, Gootenberg JS, et al. Field-deployable viral diagnostics using CRISPR-Cas13. Science 2018; 360(6387): 444–8.

38. Peterson SW, Martin I, Demczuk W, et al. Molecular Assay for Detection of Genetic Markers Associated with Decreased Susceptibility to Cephalosporins in Neisseria gonorrhoeae. J Clin Microbiol 2015; 53(7): 2042–8.

39. Shipitsyna E, Zolotoverkhaya E, Hjelmevoll SO, et al. Evaluation of six nucleic acid amplification tests used for diagnosis of Neisseria gonorrhoeae in Russia compared with an international strictly validated real-time porA pseudogene polymerase chain reaction. J Eur Acad Dermatol Venereol 2009; 23(11): 1246–53.

40. Bissessor M, Whiley DM, Fairley CK, et al. Persistence of Neisseria gonorrhoeae DNA following treatment for pharyngeal and rectal gonorrhea is influenced by antibiotic susceptibility and reinfection. Clin Infect Dis 2015; 60(4): 557–63.

41. Goire N, Lahra MM, Chen M, et al. Molecular approaches to enhance surveillance of gonococcal antimicrobial resistance. Nat Rev Microbiol 2014; 12(3): 223–9.

42. Wittwer CT. High-resolution DNA melting analysis: advancements and limitations. Hum Mutat 2009; 30(6): 857–9.

43. Chen JS, Ma E, Harrington LB, et al. CRISPR-Cas12a target binding unleashes indiscriminate single-stranded DNase activity. Science (New York, NY) 2018; 360(6387): 436–9.

44. Broughton JP, Deng X, Yu G, et al. CRISPR-Cas12-based detection of SARS-CoV-2. Nat Biotechnol 2020; 38(7): 870–4.

45. Wang S, Li H, Kou Z, et al. Highly sensitive and specific detection of hepatitis B virus DNA and drug resistance mutations utilizing the PCR-based CRISPR-Cas13a system. Clin Microbiol Infect 2021; 27(3): 443–50.

46. Donà V, Kasraian S, Lupo A, et al. Multiplex Real-Time PCR Assay with High-Resolution Melting Analysis for Characterization of Antimicrobial Resistance in Neisseria gonorrhoeae. J Clin Microbiol 2016; 54(8): 2074–81.

47. Wind CM, Bruisten SM, Schim van der Loeff MF, Dierdorp M, de Vries HJC, van Dam AP. A Case-Control Study of Molecular Epidemiology in Relation to Azithromycin Resistance in Neisseria gonorrhoeae Isolates Collected in Amsterdam, the Netherlands, between 2008 and 2015. Antimicrob Agents Chemother 2017; 61(6).

48. Ryan L, Golparian D, Fennelly N, et al. Antimicrobial resistance and molecular epidemiology using whole-genome sequencing of Neisseria gonorrhoeae in Ireland, 2014-2016: focus on extended-spectrum cephalosporins and azithromycin. Eur J Clin Microbiol Infect Dis 2018; 37(9): 1661–72.

49. Jacobsson S, Golparian D, Cole M, et al. WGS analysis and molecular resistance mechanisms of azithromycin-resistant (MIC >2 mg/L) Neisseria gonorrhoeae isolates in Europe from 2009 to 2014. J Antimicrob Chemother 2016; 71(11): 3109–16.

